# aweMAGs: a fully automated workflow for quality assessment and annotation of eukaryotic genomes from metagenomes

**DOI:** 10.1101/2023.02.08.527609

**Authors:** Davide Albanese, Claudia Coleine, Laura Selbmann, Claudio Donati

## Abstract

Metagenomics is one of the most promising approaches to identify and characterize novel microbial species from environmental samples. While a large amount of prokaryotic metagenome assembled genomes (MAGs) have been published, only a few examples of eukaryotic MAGs have been reported. This is in part due to the absence of dedicated and easy-to-use processing pipelines. Quality assessment, annotation and phylogenomic placement of eukaryotic MAGs involve the use of several computational tools and reference databases that are often difficult to collect and maintain. We present metashot/aweMAGs, a fully automated workflow capable of performing all these steps. metashot/aweMAGs can run out-of-the-box on any platform that supports Docker, Singularity and Nextflow, including computing clusters or batch systems in the cloud.

## Introduction

Metagenomics is widely used to assemble environmental and host-associated prokaryotic MAGs, helping to expand the Tree of Life with unculturable new species and providing information on the metabolic potential of members of complex microbial communities. Given the complexity of eukaryotic genomes, only few metagenomic studies include them (West *et al.*, 2018). The main obstacle is their larger and more complex genomes (Massana and López-Escardó, 2022). Moreover, the high throughput quality assessment and annotation of eukaryotic MAGs is hampered by the need to implement complex analysis pipelines, including several bioinformatic tools and maintaining multiple reference databases. EukMetaSanity (Neely et al. 2021) is a recent bioinformatic pipeline which simplifies the taxonomic classification and the gene prediction processes but it lacks the support for the assessment of genomes quality, dereplication, and phylogenetic placement.

Here, we propose metashot/aweMAGs (automated workflow for eukaryotic MAGs), an easy-to-use, container-enabled workflow for automated quality assessment, filtering, dereplication, and characterization of eukaryotic genomes and MAGs. metashot/aweMAGs can run out-of-the-box on any platform that supports Nextflow (Di Tommaso *et al.*, 2017), Docker (https://www.docker.com/) or Singularity (https://sylabs.io/singularity), including computing clusters or batch infrastructures in the cloud. The results obtained from this workflow can be used “as it is” or represent a starting point for more focused and specialized analyses on the taxa of interest.

## Workflow description

Metashot/aweMAGs is written using Nextflow (Di Tommaso *et al.*, 2017), a framework for building scalable scientific workflows using containers (Silver, 2017) allowing implicit parallelism (i.e. capability of automatically execute tasks in parallel) on a wide range of computing platforms. Reproducibility of the workflow results is guaranteed by workflow versioning (i.e. releases on GitHub) and versioned Docker images. These images enclose software tools together with their dependencies, allowing isolation from the host environment and portability across platforms. Moreover, the Nextflow workflow manager and the Docker images guarantee that the proposed pipeline can be run in a wide variety of computational environments, from local computers to high performance computing (HPC) clusters (e.g. equipped with job scheduling systems like SGE or Slurm) and cloud platforms (e.g. AWS batch service).

The workflow takes a series of genomes or metagenomic bins in FASTA format. To recover the bins, metagenomic sequence reads are first quality checked and then assembled into contigs. Among others, metaSPAdes (Nurk *et al.*, 2017) has been shown to be able to efficiently handle short and long read sequencing data and provides an experimental protocol for hybrid assembly. Finally, contigs that are likely to belong to the same organism are grouped by specific algorithms, creating the metagenomic bins (Yue *et al.*, 2020). metashot/aweMAGs v1.0.0 is composed of different modules (Figure 1) and includes several custom scripts, designed to manipulate the output of the different tasks. Required reference databases can be provided explicitly by the user; alternatively, they are automatically downloaded from the Internet. The main bioinformatic tools included and their versions are reported in Table 1. Detailed description of the command line options and parameters are available in the online documentation at https://github.com/metashot/awemags. The main steps implemented in the workflow are described as follows.

**Table 1.** Bioinformatic tools end versions included in metashot/aweMAGs v1.0.0. During the execution of the pipeline, each tool is downloaded automatically from the Internet in the form of Docker or Singularity image.

| Software | Version |
| --- | --- |
| BUSCO | 5.1.3 |
| BBTools | 38.79 |
| dRep | 2.6.2 |
| MUSCLE | 5.1 |
| RAxML | 8.2.12 |
| MMseqs2 | 13 |
| MetaEuk | 5 |
| EggNOG-mapper | 2.1.5 |
| ITSx | 1.1.2 |

**Fig. 1.**
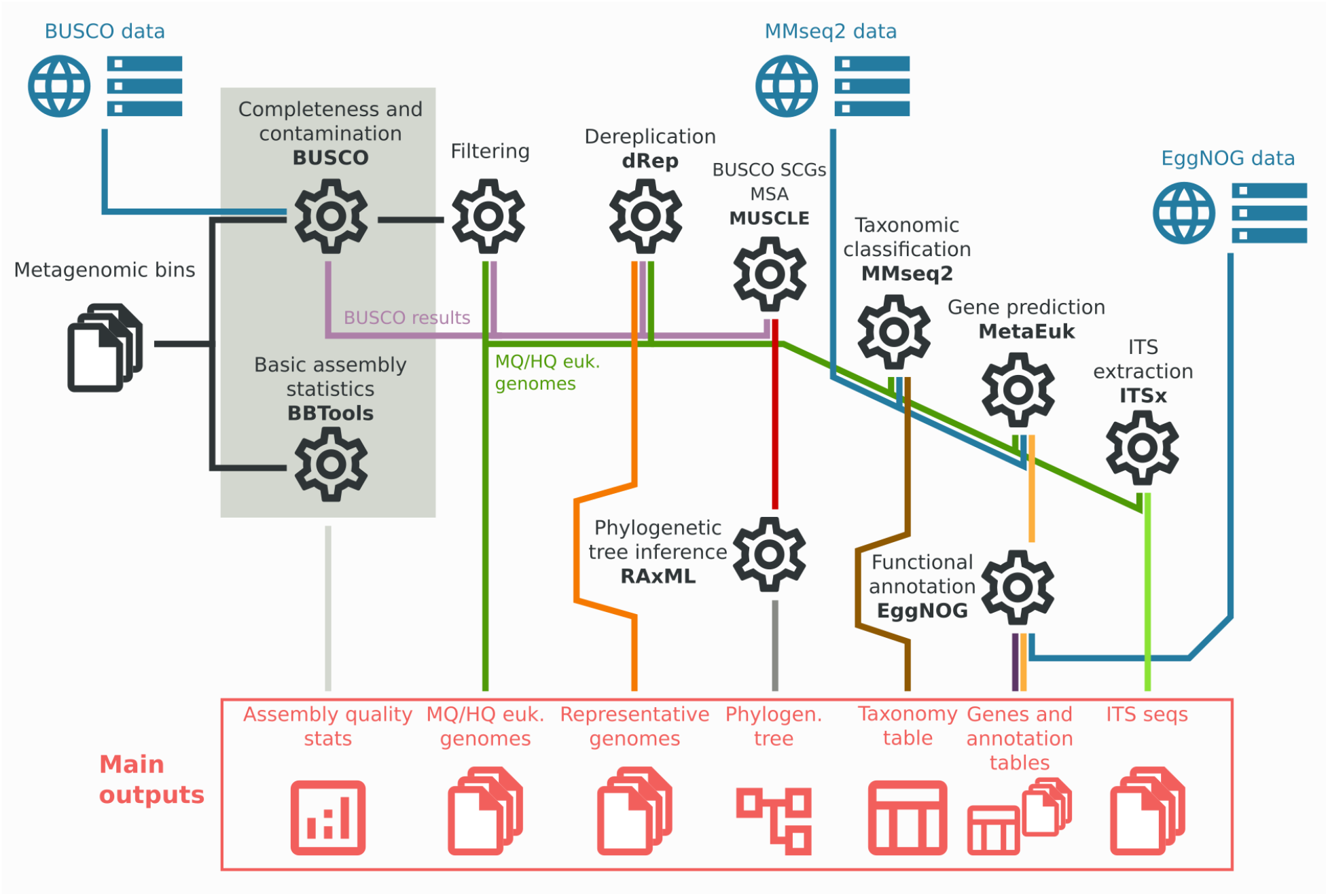
Main modules and outputs of metashot/aweMAGs v1.0.0. The workflow takes a series of genomes/metagenomic bins in FASTA format and returns: i) a tab-separated values (TSV) file including the quality information (“Assembly quality stats”) for each bin; ii) two directories, one containing the bins filtered according the completeness and contamination thresholds (i.e. medium and high quality eukaryotic genomes, in the figure reported as “MQ/HQ euk. genomes”) and the other containing representative genomes after dereplication; iii) a phylogenetic tree combined with the BUSCO’s SGC (single copy genes) multiple sequence alignments (MSA), iv) a taxonomy table, v) the predicted genes and the EggNOG’s transferred annotation tables and vi) a directory containing the predicted internal transcribed spacer ITS sequences.

### Quality assessment, filtering and dereplication

Quality assessment is performed for each input genome/metagenomic bin using BUSCO (Manni *et al.*, 2021) v5 and the BBTools’s (https://jgi.doe.gov/data-and-tools/software-tools/bbtools/); “statswrapper” program. The workflow reports in a single tab-separated values (TSV) file the estimated completeness and contamination of each bin, as well basic assembly statistics, namely the genome size, the number of contigs and the N50/L50. metashot/aweMAGs supports the BUSCO’s automated lineage selection, through which the optimal domain for each input bin is selected. Alternatively, the user can even explicitly choose between the different BUSCO lineage datasets.

The input metagenomic bins are (optionally) filtered according to the completeness and contamination thresholds (by default, 50% and 10% for completeness and contamination, respectively). The bins passing these thresholds (henceforth called MAGs) are placed in a specific folder.

Dereplication is a procedure that clusters the input genomes according to their genome similarity, using measures such as the Average Nucleotide Identity (Goris *et al.*, 2007) (ANI). Dereplication is used in metagenomic context to account for small differences between closely related genomes, possibly related to assembly artifacts. For prokaryotes, a threshold of 95% ANI has been shown to define species-level clusters (Jain *et al.*, 2018). The workflow optionally clusters the input genomes using dRep (Olm *et al.*, 2017) (default ANI threshold 99%). For each cluster, the genome with the higher dRep score is selected as representative. In case the filtering procedure was performed, the score is computed using the formula s = c_m_ - 5 x c_n_ + 0.5 x log(N50), where c_m_ and c_n_ are the completeness and contamination, respectively; otherwise, the score is computed as s = log(g_s_), where g_s_ is the genome size.

### BUSCO SGC multiple sequence alignment and phylogenetic placement

For each BUSCO single-copy gene (SCG) a multiple sequence alignment (MSA) is carried out using MUSCLE v5 (Edgar, 2022). Performing this step is not possible when the BUSCO automated lineage selection is activated. This is due to the fact that BUSCO might select different SGC datasets for each input.

For each SCG MSA, columns represented in <50% of the input genomes or columns with less than 25% or more than 95% amino acid consensus are trimmed in order to remove sites with weak phylogenetic signals (Rinke *et al.*, 2021). To reduce the total number of columns selected for tree inference, the alignment was further trimmed by randomly selecting ⎣p_c_ / n_SGC_⎦ columns, where p_c_ is the user parameter that specifies the maximum number of columns for the final MSA (default 5000) and n_SGC_ is the total number of BUSCO SGC for the specified lineage. The trimmed MSAs are then concatenated into a single MSA and the phylogenomic tree inferred using RAxML. Two are the modes available for RAxML:

- default mode: it constructs a maximum likelihood (ML) tree. This mode runs the default RAxML tree search algorithm (Stamatakis *et al.*, 2007) and performs multiple searches for the best tree (10 distinct randomized MP trees by default).
- rbs mode: it assesses the robustness of inference and constructs a ML tree. This mode runs the rapid bootstrapping full analysis (Stamatakis, Hoover and Rougemont, 2008). The bootstrap convergence criterion or the number of bootstrap searches can be specified.

### Taxonomic classification and ITS detection

Taxonomic annotation is delegated to the MMseqs2 easy-taxonomy workflow (Mirdita *et al.*, 2021). This workflow is able to perform fast taxonomic assignments to metagenomic contigs extracting all possible protein fragments and searching them against a reference database. Since MMseqs2 easy-taxonomy works on contiguous sequences, before the classification each genome contig is concatenated into a single pseudochromosome using the sequence “NNNNNCATTCCATTCATTAATTAATTAATGAATGAATGNNNNN” as separator. This sequence provides a stop codon and a start site in all six reading frames (Tettelin *et al.*, 2005), guaranteeing that chimera genes spanning different contigs are not created by concatenating the sequences. As mentioned before, this step requires a MMseqs2 protein database (https://github.com/soedinglab/mmseqs2/wiki#downloading-databases) augmented with taxonomic information, which can be provided by the user or downloaded automatically from the Internet.

For eukaryotes, and especially for fungi, the ITS region still has an important role in species identification and phylogenetic inference (Eberhardt, 2010; China Plant BOL Group *et al.*, 2011). Using the software ITSx (Bengtsson-Palme *et al.*, 2013), the proposed workflow optionally extracts, for each input MAG, the full-length ITS sequences, including the ITS1, 5.8S and ITS2 subregions.

### Gene prediction and functional annotation

Gene prediction is performed using the MetaEuk easy-predict software. MetaEuk is a reference-based tool for gene discovery in eukaryotic contigs with good sensitivity and speed (Karin, Mirdita and Söding, 2020). Also in this case, this step requires a reference MMseqs2 database.

The translated coding sequences (CDS) predicted by MetaEuk are functionally annotated using EggNOG-mapper (Cantalapiedra *et al.*, 2021). Annotation is performed searching against the precomputed orthologous groups from the eggNOG database (Huerta-Cepas *et al.*, 2019) v5.0. The database integrates functional annotations collected from several sources, including Gene Ontology (Harris *et al.*, 2004) (GO) terms, Kyoto Encyclopedia of Genes and Genomes (KEGG) functional orthologs (Kanehisa *et al.*, 2017) and Clusters of Orthologous Groups (COG) (Tatusov *et al.*, 2000) categories. For each transferred annotation, the workflow reports a TSV file, including the feature counts for each input MAG.

## Hardware requirements

As the different computational steps involve softwares with widely varying computational requirements, the infrastructure needs to be adequate for performing each step separately. However, since the requirements of some of these steps (e.g. MSA) vary widely depending on the input data, we have implemented an adaptive strategy taking advantage of the retry-if-fail feature of the Nextflow workflow manager. In this approach, the hardware requirements for a particular process are progressively increased if the step fails due to insufficient resources. This is done by informing in a transparent way the job executor used (e.g. the queue or the batch system). This strategy proved to be very effective in increasing the efficiency of the workflow, avoiding unnecessary overbooking of resources (Albanese and Donati, 2021).

The most critical task in terms of memory usage is the reference-based gene discovery, due to the MetaEuk v5 requirements. Therefore, metashot/aweMAGs v1.0.0 requires a minimum of 16GB of RAM when a small MMseqs2 database like the Swiss-Prot is used and more than 100GB of RAM when a relatively big genome (e.g. 1 Gbp-long) is analyzed using the UniRef50 database.

All the system-specific configurations are specified by the user in a single configuration file. An example is provided at the metashot webpage https://metashot.github.io/#dependencies.

## Full workflow run

In this section, we show how to perform the full workflow on a set of input metagenomic bins. Binning (Yue *et al.*, 2020) is the procedure that usually follows the metagenomic assembly or co-assembly, and is needed for grouping contigs that are likely to belong to the same organism, increasing the interpretability of metagenomic data. As mentioned previously, the software prerequisites for running metashot/aweMAGs on a POSIX compatible system are Nextflow and Docker (or Singularity). Given a series of candidate metagenomic bins in FASTA format stored in the “bins” directory, the version 1.0.0 of the workflow can be run with the following command line:

nextflow run metashot/awemags -r 1.0.0 --genomes ‘bins/*.fa’

This command runs the full pipeline with the BUSCO auto lineage mode for the eukaryotes (default). The reference databases needed for the different analysis (i.e. BUSCO, MMseqs2 and eggNOG) are downloaded automatically from the Internet and the results are stored in the default directory “results”. For the documentation and the complete list of options see the GitHub page https://github.com/metashot/awemags.

## Extracting and annotating medium and high quality eukaryotic plankton genomes from Tara Oceans SMAGs

In a recent paper (Delmont *et al.*, 2022), Delmont et al. report 683 eukaryotic MAGs and 30 single amplified genomes (SAGs) from marine plankton, providing an important resource to interpret marine eukaryotic diversity. Briefly, starting from 798 metagenomes and 158 eukaryotic single cells sequences, the authors performed assembly, manually curated binning and dereplication of the genomes, obtaining a non-redundant database of 713 eukaryotic genomes, the “SMAGs”. The taxonomic classification of the SMAGs was determined using a combination of five approaches and the support of the Marine Eukaryote Transcriptomes reference database (Niang *et al.*, 2020) (METdb). Moreover, the authors predicted the protein coding regions for each SMAG using three complementary approaches (protein alignments against reference databases, metatranscriptomic assemblies mapping and *ab-initio* predictions).

Here, to demonstrate the potentiality of metashot/aweMAGs, we filtered and re-analyzed the SMAGs in an automatic manner. The SMAGs were downloaded from the website https://www.genoscope.cns.fr/tara/ and analyzed using metashot/aweMAGs v1.0.0 on a Sun Grid Engine (SGE) batch-queuing system (version 8.1.9). Since we wanted to produce a phylogenetic analysis on the entire eukaryotic domain, we set the BUSCO lineage parameter (“--lineage”) to “eukaryota” (version ODB10). In this way, the BUSCO set of 255 eukaryotic single-copy core gene markers was recovered for each input SMAG. For the taxonomic classification and gene prediction processes, the MMseqs2 UniRef50 (Suzek *et al.*, 2007) (release 2022_01) was used as the reference protein database. Functional annotation was performed using the EggNOG mapper database version 5.0.2. The analysis required a total of 15.615 CPU-hours, with most of the time used for the reference-based gene discovery (with a mean of 19 hours per sample with 8 CPUs).

The workflow selected and annotated a total of 176 between medium quality (MQ, completeness >50%, contamination <10%) and high quality (HQ, completeness >90%, contamination <5%) SMAGs, including the 1.32 Gbp-long Bacillariophyta genome *(Delmont et al., 2022)* (see Supplementary Table S1). The rooted maximum likelihood (ML) phylogenetic tree based on the 255 BUSCO eukaryotic single-copy gene markers is shown in Figure 2a.

**Fig 2.**
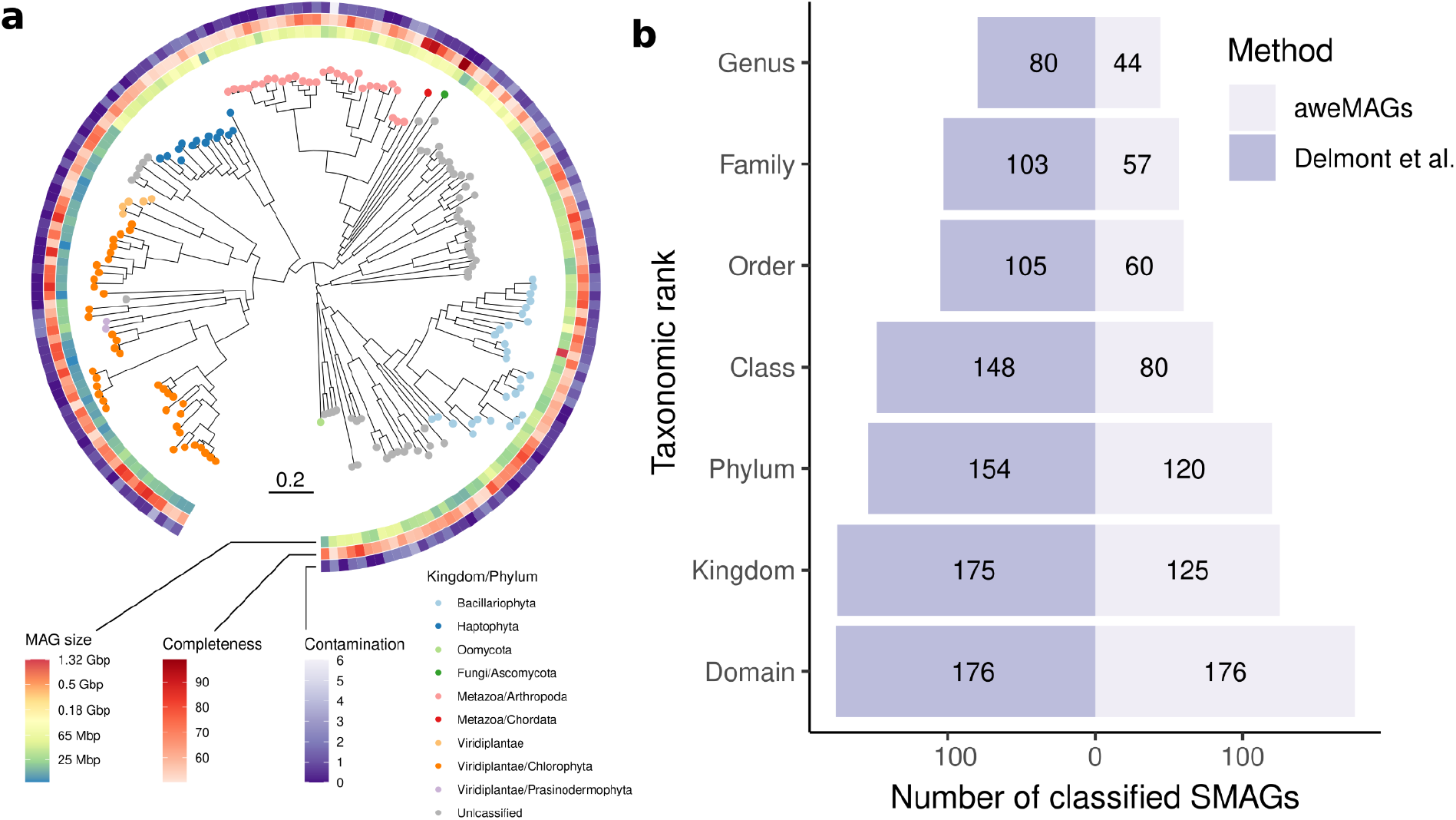
MQ and HQ SMAGs from Delmont and colleagues. a) Rooted ML phylogenetic tree representing the 176 MQ e HQ eukaryotic SMAGs based on the 255 BUSCO eukaryotic single-copy gene markers b) Number of taxonomically classified SMAGs in (Delmont *et al.*, 2022) and using the workflow presented in this work (right). Marine stramenopile (MAST) lineages classifications are not considered in this figure.

The taxonomic classification procedure implemented in metashot/aweMAGs recognized kingdom end genera in 125 and 44 SMAGs, respectively. Although the manually curated procedure described in (Delmont *et al.*, 2022) allowed to identify the SMAGs at deeper levels (Figure 2b), we found no inconsistencies between the two methods (see Supplementary Table S2), with the exception of the SMAG TARA_PSE_93_MAG_00199, classified by our workflow as Leotiomycetes instead of Ascomycetes (class rank). However, this discrepancy is probably related to the different standards used for taxonomy, as the class Leotiomycetes is present in the reference taxonomy database used by the MMseqs2 easy-taxonomy workflow that is included in our pipeline, while Ascomycetes is only referred to all fungi in the phylum Ascomycota (Wijayawardene *et al.*, 2017). Many of the 51 unclassified genomes at the kingdom rank are classified in (Delmont *et al.*, 2022) as Chromista (n=39) and 19 are in MAST (marine stramenopiles) lineages (Massana *et al.*, 2004, 2014). We predicted and annotated a total of 2,609,089 CDS. The proposed workflow exports a series of annotation profiles including the KEGG orthologs (ko), modules and pathways. The annotation matrix for the KEGG orthologs is included in the Supplementary Table S3.

## Discussion

In this paper, we present aweMAGs, a software pipeline for the quality assessment, taxonomic classification and functional annotation of metagenome assembled genomes for Eukaryotes. While MAGs are routinely assembled and characterized for Prokaryotes, the study of the eukaryotic component of the microbiota is still in its infancy. One of the major obstacles, hampering the research, is the lack of easy-to-use and portable computational pipelines that can analyze a wide range of datasets and provide data, which can be compared between different projects. For this reason, we developed aweMAGS, a portable software pipeline implemented using widely used software technologies such as Nextflow, Docker and Singularity that guarantee portability across a wide range of computational infrastructures.

The pipeline has been tested on one recent dataset of eukaryotic MAGs and SAGs that were used as a gold standard for taxonomic classification. The results obtained using our pipeline were fully consistent with the results provided by (Delmont *et al.*, 2022), showing that an automated pipeline could, without human intervention, provide results that are consistent with a careful, complex and labor intensive analysis that manually integrates different computational approaches.

However, a fully automated approach like the one implemented in aweMAGs has limitations. A manual refinement, the use of complementary methods and specialized databases would allow to obtain more accurate results (e.g for gene prediction) or with higher resolution, as in the case of taxonomic classification in comparison with the original work by (Delmont et al. 2022). However, manual interventions are difficult to document, often leading to poor repeatability of the bioinformatic analyses and thus hampering the comparison of results across different studies. In addition, manual curation of large datasets poses significant problems in terms of consistency, especially if a team of curators is involved. For these reasons, we believe that the aweMAGs is a useful tool to support the rapid implementation of comparative metagenomic studies, representing a starting point for more specialized analyses and reducing the burden of manual curation by human experts only to those cases when a deeper analysis is needed.

## Software availability

Software and documentation are freely available at https://github.com/metashot/awemags under the GNU General Public License v3.0. The workflow release used in this work (v1.0.0) is available at https://github.com/metashot/awemags/releases/tag/1.0.0. The prebuilt Docker images are downloadable from the Docker registry https://hub.docker.com/u/metashot and their definitions are available at https://github.com/metashot/docker.

